# *Cis*-regulatory variants affect gene expression dynamics in yeast

**DOI:** 10.1101/2021.03.16.435665

**Authors:** Ching-Hua Shih, Justin C. Fay

## Abstract

Evolution of *cis*-regulatory sequences depends on how they effect gene expression and motivates both the identification and prediction of *cis*-regulatory variants responsible for expression differences within and between species. While much progress has been made in relating *cis*-regulatory variants to expression levels, the timing of gene activation and repression may also be important to the evolution of *cis*-regulatory sequences. We investigated allele-specific expression (ASE) dynamics within and between *Saccharomyces* species during the diauxic shift and found appreciable *cis*-acting variation in gene expression dynamics. Within species ASE is associated with intergenic variants, but ASE dynamics are more strongly associated with insertions and deletions than ASE levels. To refine these associations we used a high-throughput reporter assay to test promoter regions and individual variants. Within the subset of regions that recapitulated endogenous expression we identified and characterized *cis*-regulatory variants that affect expression dynamics. Between species, chimeric promoter regions generate novel patterns and indicate constraints on the evolution of gene expression dynamics. We conclude that changes in *cis*-regulatory sequences can tune gene expression dynamics and that the interplay between expression dynamics and other aspects expression are relevant to the evolution of *cis*-regulatory sequences.

## Introduction

*Cis*-regulatory sequences control the activation and repression of genes and are thought to play a central role in evolution (Carroll, 2005). However, genetic analysis of phenotypic variation frequently uncovers changes in protein coding sequences (Stern and Orgogozo, 2008; Fay, 2013). One limitation of genetic mapping and transgenic studies is identifying small effect mutations. If evolution predominantly occurs through numerous changes of small effect (Rockman, 2012), the role of *cis*-regulatory sequences is harder to discern. Even so, *cis*-regulatory changes have been shown to be important in polygenic adaptation (Bullard et al., 2010; Fraser et al., 2010, 2011, 2012; Naranjo et al., 2015), the accumulation of multiple changes at evolutionary hotspots (Frankel et al., 2011; Engle and Fay, 2012, 1; Martin and Orgogozo, 2013; Li and Fay, 2019), and variation in fitness and disease (Boyle et al., 2017; Sharon et al., 2018). Regardless of the overall role of *cis*-regulatory sequences, understanding how variation in *cis*-regulatory sequences generates variation in gene expression is important to understanding the evolution of gene regulation.

Across organisms there is an abundance of *cis*-acting sequence variation that affects gene expression levels (Hill et al., 2020). The causes of *cis*-regulatory variation are not as easily characterized. Transcription factor binding sites are often found to play important roles (Zheng et al., 2011). However, the number, identity and position of binding sites can also vary without affecting expression. The flexibility of *cis*-regulatory sequences is shown by genes with similar expression patterns but different *cis*-regulatory sequences (Berman et al., 2002). In the case of orthologous genes from different species, binding site turnover and transcription factor re-wiring explain substantial divergence in *cis*-regulatory sequences without expression divergence (Ludwig et al., 2000; Dermitzakis and Clark, 2002; Hare et al., 2008; Tuch et al., 2008; Venkataram and Fay, 2010; Swanson et al., 2011; Bergen et al., 2016). Consequently, predicting changes in gene expression based on variation in individual transcription factor binding sites has proven difficult (Doniger and Fay, 2007; Doniger et al., 2008).

Despite the flexibility of binding sites within *cis*-regulatory sequences, sequences flanking binding sites evolve under constraints and can affect expression. For example, over a third of yeast intergenic sequences are estimated to be under selective constraint (Chin et al., 2005; Doniger et al., 2005). This fraction is greater than that expected from either conserved or experimentally identified binding sites (Doniger et al., 2005; Venkataram and Fay, 2010). Sequences flanking binding sites have also been shown to affect expression, potentially related to DNA shape, nucleosome positioning or weak binding sites (Tanay et al., 2005; White et al., 2013; Abe et al., 2015; Levo et al., 2015; Inukai et al., 2017). Consequently, conservation scores have proven important for predicting *cis*-regulatory variants (Huang et al., 2017; Kircher et al., 2019; Renganaath et al., 2020).

*Cis*-regulatory sequences can affect other aspects of gene regulation besides expression levels. Stochastic noise in gene expression provides a mechanism for bet-hedging strategies (Raj and van Oudenaarden, 2008) and is encoded by and evolves through changes in *cis*-regulatory sequences (Richard and Yvert, 2014). *Cis*-regulatory variants that alter noise in expression levels have been shown to be under selection and can occur both within and outside of known binding sites (Carey et al., 2013; Sharon et al., 2014; Metzger et al., 2015; Schor et al., 2017; Duveau et al., 2018).

Gene expression dynamics, which include the timing and rate of gene activation and repression, are also important aspects of gene regulation (López-Maury et al., 2008; Yosef and Regev, 2011). Gene expression dynamics can be altered by transcription factors and their interactions with promoters, but also depend on nucleosomes and their positions relative to binding sites (Lam et al., 2008; Hager et al., 2009; Dadiani et al., 2013; Hansen and O’Shea, 2015). Notably, chromatin mutants slow gene activation without compromising final levels of gene expression (Barbaric et al., 2001; Floer et al., 2010). Variation in *cis*-regulatory sequences can also affect gene expression dynamics but these dynamics are only sometimes captured (Ackermann et al., 2013; Francesconi and Lehner, 2014; Strober et al., 2019). Thus, the causes of *cis*-regulatory variation in gene expression dynamics have not been characterized nor their relationship to variation in gene expression levels.

In this study we investigate *cis*-acting variation in gene expression dynamics. We survey and find allele-specific differences in expression dynamics both within and between *Saccharomyces* species during the diauxic shift, when there is major transition from the expression of genes involved in fermentation to respiration (DeRisi et al., 1997). Using these data, we associated allele-specific expression (ASE) with promoter variation and individual variants using a high-throughput reporter assay. Our results inform our understanding of variation in gene expression dynamics and point towards an integrated view of gene expression and how it evolves.

## Results

### *Cis*-regulatory variation in gene expression levels and dynamics

To identify *cis*-regulatory variation in gene expression dynamics we measured allele-specific expression (ASE) in three intra-specific and two inter-specific diploid hybrids. Hybrids were generated by crossing a North American *S. cerevisiae* strain (Oak) to an *S. cerevisiae* wine strain (Wine) and two strains from China (China I and China II), as well as to a strain of *S. paradoxus* and *S. uvarum* (Table S1), enabling us to examine a range of divergence in gene regulation. To capture temporal differences in ASE that occur during the diauxic shift we generated RNA-sequencing data from 19 time-points for each hybrid, spanning the shift from fermentation to respiration as measured by glucose depletion (Figure S1).

Allele-specific expression requires RNA-sequencing reads that can be distinguished as coming from one of the two parental strains. To measure ASE while avoiding mapping bias (Degner et al., 2009; Stevenson et al., 2013), we mapped reads to the combined parental genomes and enumerate allele-specific reads. The proportion of reads mapping to each parental genome was equivalent across all 5 hybrids, except for one arm of chromosome XIII in the China I hybrid consistent with aneuploidy (Figure S2), which we removed from subsequent analysis.

Two statistical tests were used to separately identify genes exhibiting differences in ASE levels and changes in ASE dynamics over time. Genes with ASE dynamics were identified by testing for an autocorrelation in the ratio of allele-specific reads over time. Under the null model the ratio of the two alleles is constant over time but not necessarily equal to one. Genes with ASE levels were identified by testing for differences between the expression of the two alleles across all time-points. Using these tests we found more genes showed ASE levels compared to ASE dynamics and the number of genes with either ASE levels or dynamics increased with divergence (FDR < 0.01, Table 1). As expected, genes with ASE levels showed larger average allele differences across time-points, and genes with ASE dynamics showed larger standard deviations in allele differences across time-points (Figure S3). Genes with ASE levels and genes with ASE dynamics were relatively evenly distributed across genes whose expression increased/decreased or showed a peak/trough during the diauxic shift (Figure S4, Table S2).

**Table 1.** Number of genes with allele-specific expression

| Group<sup>1</sup> | Intra-specific hybrids<sup>3</sup> |  |  | Inter-specific hybrids<sup>3</sup> |  |
| --- | --- | --- | --- | --- | --- |
|  | *S. cerevisiae*<br>(Oak x Wine) | *S. cerevisiae*<br>(Oak x China II) | *S. cerevisiae*<br>(Oak x China I) | *S. cerevisiae*<br>x<br>*S. paradoxus* | *S. cerevisiae*<br>x<br>*S. uvarum* |
|  | YJF1460 | YJF1455 | YJF1454<sup>2</sup> | YJF1453 | YJF1484 |
| Dynamics | 671 | 699 | 911 | 2055 | 1827 |
| Both | 371 | 375 | 501 | 1260 | 1253 |
| Levels | 1964 | 2088 | 2260 | 2930 | 3237 |

<sup>1</sup>Group: genes with significant (FDR < 0.01) allele-specific differences in dynamics, levels, or both dynamics and levels.
<sup>2</sup>The total number of genes is 4703 except for 358 genes on chromosome 13R of the China I hybrid that were removed.
<sup>3</sup>Oak is most closely related to the Wine strain, followed by China II, China I, *S. paradoxus* and *S. uvarum*.

Changes in ASE over time can result from a variety of differences in the dynamics of the two alleles. To illustrate this variety we consider a gene that is activated during the diauxic shift (Figure 1A and 1B). ASE can be condition-specific due to an allele difference in the present but not absence of glucose, or vice-versa. ASE can also differ specifically during the diauxic shift due to a difference in the timing or rate of gene activation that does not require ASE differences before or after the shift. To characterize ASE we applied k-means clustering to ASE allele frequencies and found two types of patterns (Figure 1C, Table S3). The majority of genes (72%) showed environment-dependent ASE and the remaining genes showed an ASE maximum or minimum during the transition (Clusters 6, 9, 10, 12, Figure 1C).

**Figure 1.**
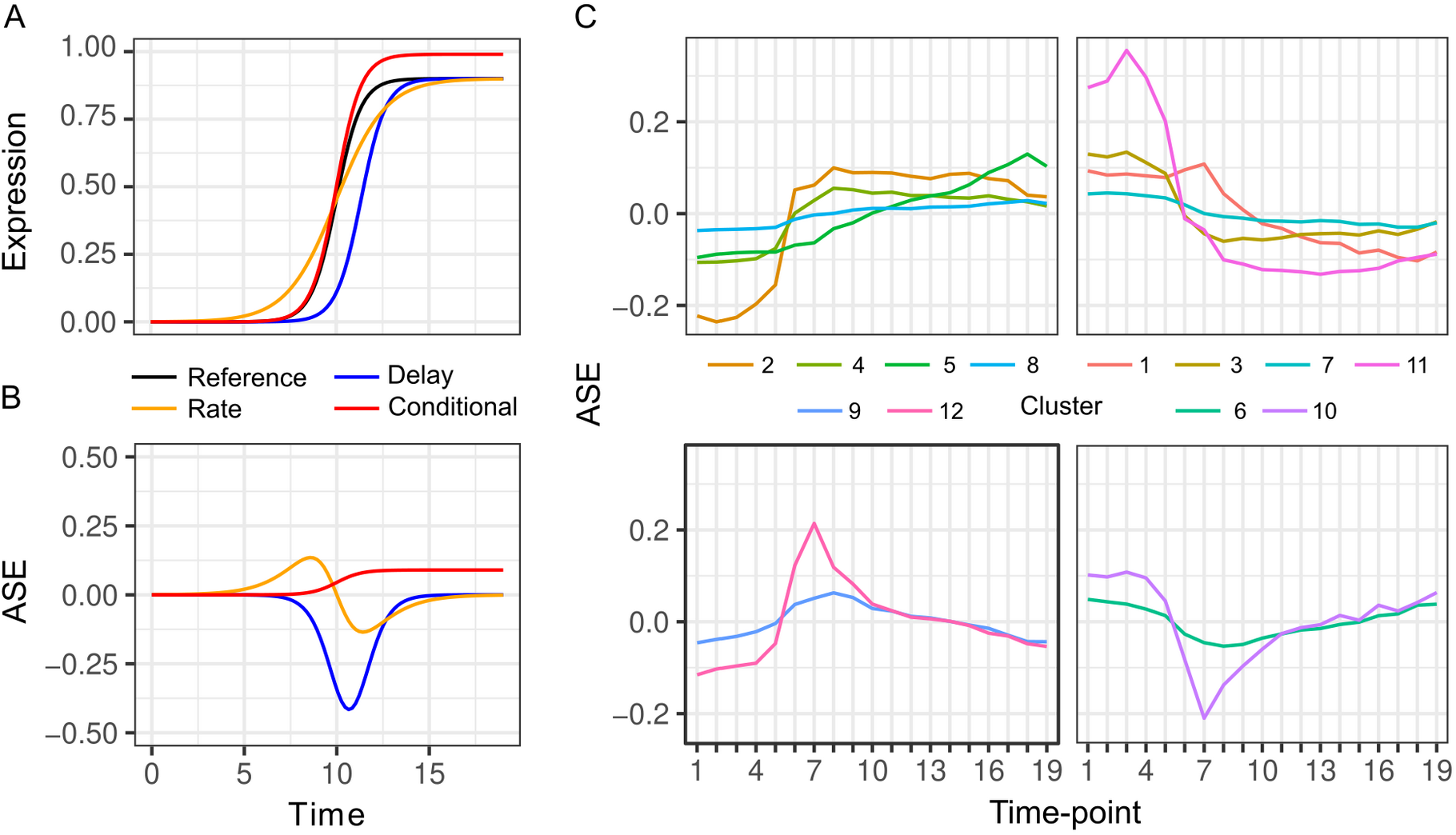
Patterns of allele-specific expression dynamics. (A) Three hypothetical types of differences in expression dynamics in comparison to a common reference (black) are shown by a time-delay (blue), rate change (orange), and condition-specific expression difference (red). (B) ASE based on the three types of differences in comparison to the common reference. (C) Average ASE across 19 time-points of k-means clustering of 6,135 genes with significant ASE dynamics. Clusters 6, 9, 10 and 12 (bottom panels) show maximum deviation during the diauxic shift, whereas the others generally show increasing or decreasing ASE differences overtime consistent with condition-specific ASE.

### ASE is associated with SNPs and InDels

ASE is caused by *cis*-acting single nucleotide polymorphisms (SNPs) or insertion deletion polymorphisms (InDels) that affects gene expression. In yeast, *cis*-acting variants most likely occur within the small (~500 bp) intergenic region upstream of a gene, but could also occur within the coding or 3’ region of a gene. For each hybrid we tested whether the number of variants in these regions predicts significant ASE levels or ASE dynamics using logistic regression.

Both ASE levels and dynamics were associated with the number of SNP and InDel variants, but these associations varied by hybrid, the type of variant, and where the variants occurred. Within intra-specific hybrids, upstream SNPs and InDels were associated with both ASE levels and dynamics, but InDels showed stronger associations as measured by the odds ratio and the significance (Figure 2). SNPs and InDels within coding and downstream regions were also associated with ASE, but the associations were weaker and/or less significant than with upstream variants (Table S4).

**Figure 2.**
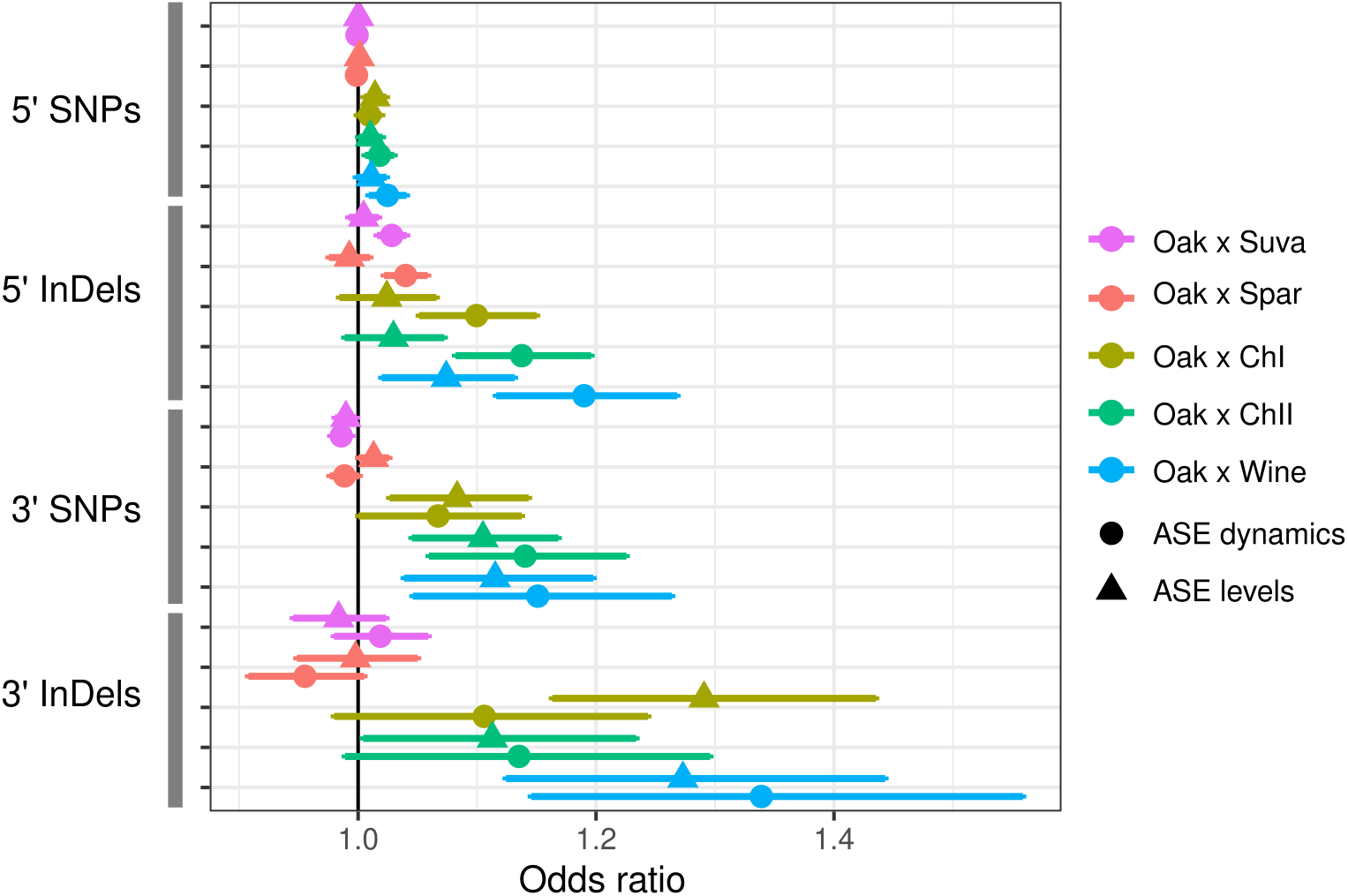
ASE is associated with intergenic SNPs and InDels. The odds ratio (OR) and 95% confidence interval for associations between the number of SNPs or InDels and significant ASE levels (triangles) and dynamics (circles). The OR of each hybrid is shown separately for upstream (5’) and downstream (3’) intergenic

Inter-specific hybrids only showed significant associations between ASE dynamics and upstream SNPs and InDels. Association between ASE and divergence may be weak or absent if most substitutions between species do not affect gene expression. The inter-specific hybrids have much higher rates of divergence compared to the intra-specific hybrids: an average of 15 and 107 upstream SNPs and InDels, respectively, compared to 1.2 and 6.0 SNPs and InDels within the intra-specific hybrids (Table S5). Consistent with the weaker inter-specific associations, the most closely related intra-specific hybrid (Oak × Wine) showed the strongest associations.

### Intra-specific *cis*-regulatory variants

The associations between ASE and the number of upstream intergenic variants indicate that promoter polymorphism is a significant contributor to ASE. To specifically measure the effects of promoter polymorphism and identify causal variants we used a high-throughput *<u>c</u>is*-<u>r</u>egulatory <u>e</u>lement (CRE-seq) reporter assay (Mogno et al., 2013). In this assay, promoter sequences are synthesized and the resulting pooled library is cloned and integrated into a single site in the yeast genome (Figure S5). The synthesized sequences include a 10 bp barcode that can be used as a tag to measure gene expression through RNA-sequencing and relative abundance through DNA-sequencing.

We designed a CRE-seq library to test promoter variants upstream of 69 genes that exhibited ASE levels and/or dynamics in the Oak × ChII hybrid. Because the synthesized promoters were limited to 130 bp, we designed five overlapping CRE sequences per gene to test all variants within the 250 bp region upstream of the transcription start site. There were a total of 337 variants, an average of 4.2 SNPs and 0.72 InDels per gene. For any CRE sequences with more than a single difference between the Oak and ChII alleles, we also generated CREs for each ChII variant in the Oak allele and vice-versa (Figure S5). The total library contained 1,818 CREs with four barcode replicates per CRE.

We first tested whether the synthetic promoter regions could recapitulate expression of the endogenous genes. We measured CRE expression over 19 time-points during the diauxic shift in the same hybrid background as RNA-seq. Out of 334 regions, 137 were correlated with RNA-seq expression (FDR < 0.05) and 17 genes had no region correlated. CRE-seq regions correlated with RNA-seq tended to lie further away from the transcription start site, and in some cases showed a gradual increase in correspondence (Figure 3). We also barcoded and measured expression from full length promoters of three genes (Figure S6). All three genes showed good correspondence between the full length promoter and a shorter CRE-seq region.

**Figure 3.**
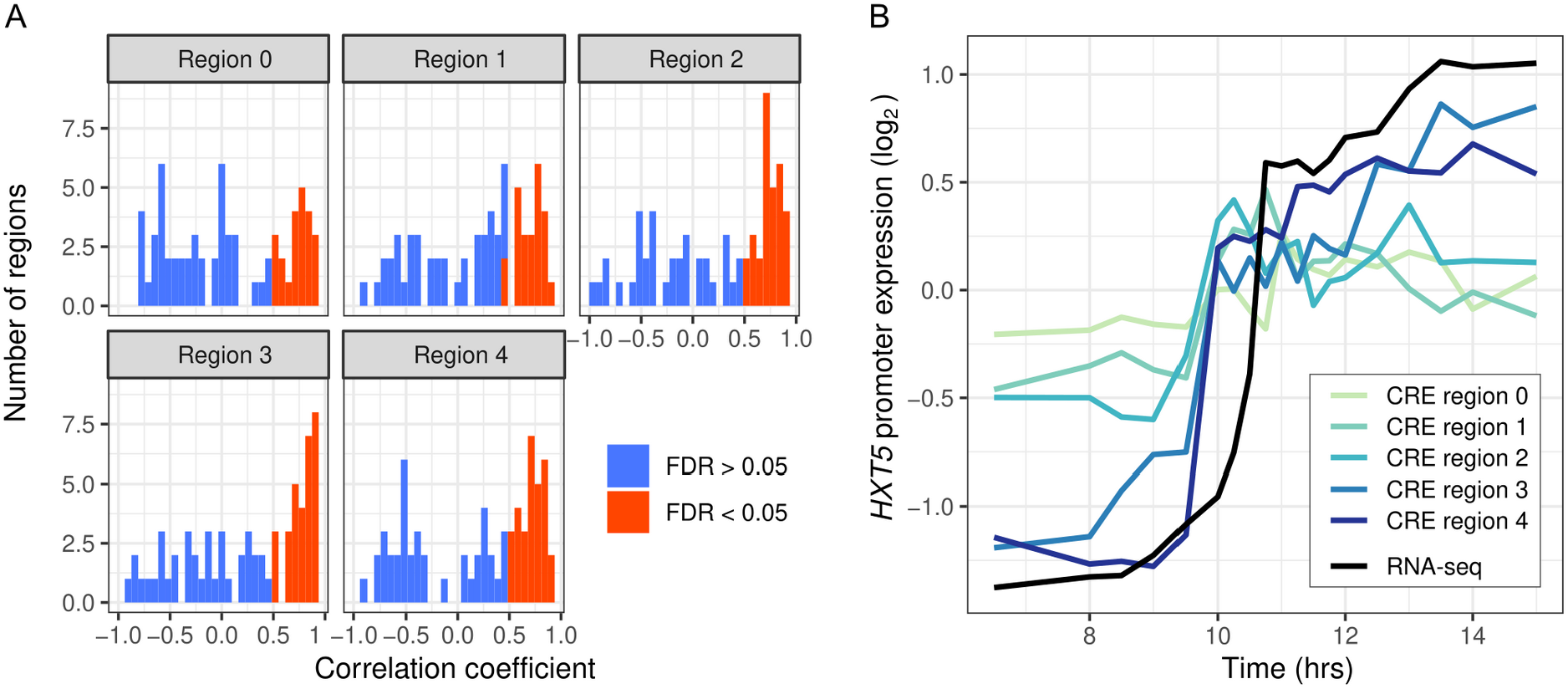
Intra-specific CREs recapitulate endogenous expression dynamics. (A) Histogram of the correlations between CRE expression and the endogenous (RNA-seq) expression of 69 genes. Correlations are shown separately for CRE regions 0-4, ordered proximal to distal of the transcription start site with shifts of 30 bp. CREs with a significant (FDR < 0.05) correlation are shown in red, the rest in blue. (B) CRE expression of five regions upstream of *HXT5* as well as its endogenous expression from RNA-seq.

However, one of the genes, *ALD5*, showed both short and full length reporter expression notably different from the endogenous RNA-seq expression. One explanation for this difference is that we used an annotated transcription start site 515 bp upstream of the ATG whereas some studies indicate a site much closer (78-84 bp) to the ATG (Zhang and Dietrich, 2005; Pelechano et al., 2013).

To determine whether any of the CRE regions contained variants that affect expression we examined the 281/334 regions upstream of the 69 genes with one or more differences between the Oak and ChII alleles. One of the regions showed significant differences in expression levels between the Oak and ChII alleles, and 31 showed differences in expression dynamics (FDR < 0.05, Table 2). Eleven of these regions had only a single variant that differentiated the two parental alleles and the rest had between 2 and 9 variants. For regions with multiple variants we tested each using CREs containing the Oak variant in the ChII background and vice-versa. For expression levels we found significant effects for one of the two variants tested, and for expression dynamics we found 35 out of 70 variants (FDR < 0.05, Table 2), four of which were detected by multiple regions.

**Table 2.**
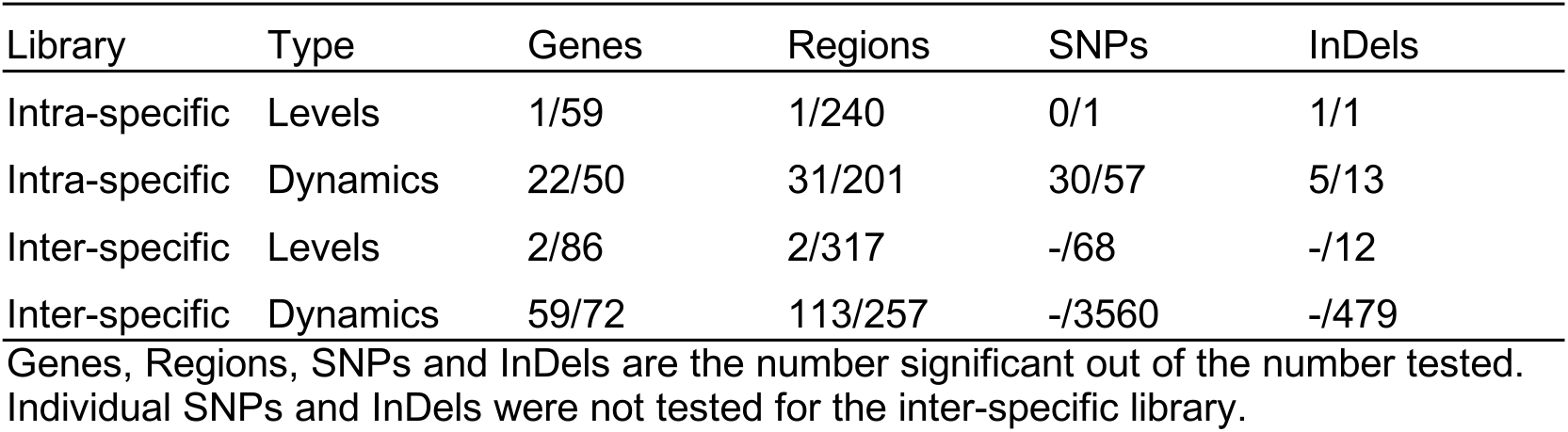
CRE regions and variants affecting gene expression

Promoter regions that showed allele-specific differences in expression dynamics had high rates of polymorphism and often multiple variants that affected expression. The rate of variants in the 31 CRE regions with differences in expression dynamics (2.1%) was higher than that of intergenic regions across the genome (1.4%)(Fisher’s exact test, P=0.0038). Out of the 22 genes with one or more CRE regions showing differences in expression dynamics between the Oak and ChII alleles, 8 had two or more significant variants, 12 had only a single variant, and two had no significant variants. While the number of InDels with significant effects on expression dynamics was small, the ratio of significant SNPs to InDels (6.0) was not different from that of intergenic regions across the genome (4.8)(Fisher’s exact test, P > 0.05).

Variants with significant effects on CRE-seq expression were not always consistent with patterns of RNA-seq ASE (Figure 4). Only 16 of the 31 regions showing CRE-seq expression dynamics correlated with RNA-seq expression. For example, region 4 of the *YPS6* promoter showed increased expression over time in the RNA-seq data but not in the CRE-seq data. Even so, the Oak allele exhibited higher expression levels than the ChII allele in both the RNA-seq and CRE-seq assays, a difference that can be attributed to one of the two variants in the region. The *ICL2* promoter showed increased expression over time in both the RNA-seq and CRE-seq assays, but not all CRE-seq allele differences were consistent with the RNA-seq allele differences. Region 1 of the *ICL2* promoter showed allele-specific expression differences consistent with RNA-seq and multiple variants with effects on expression dynamics. However, region 3, which had only a single variant, showed allele differences in expression dynamics inconsistent with RNA-seq, whereby the Oak allele responded more strongly than the ChII allele to the diauxic shift.

**Figure 4.**
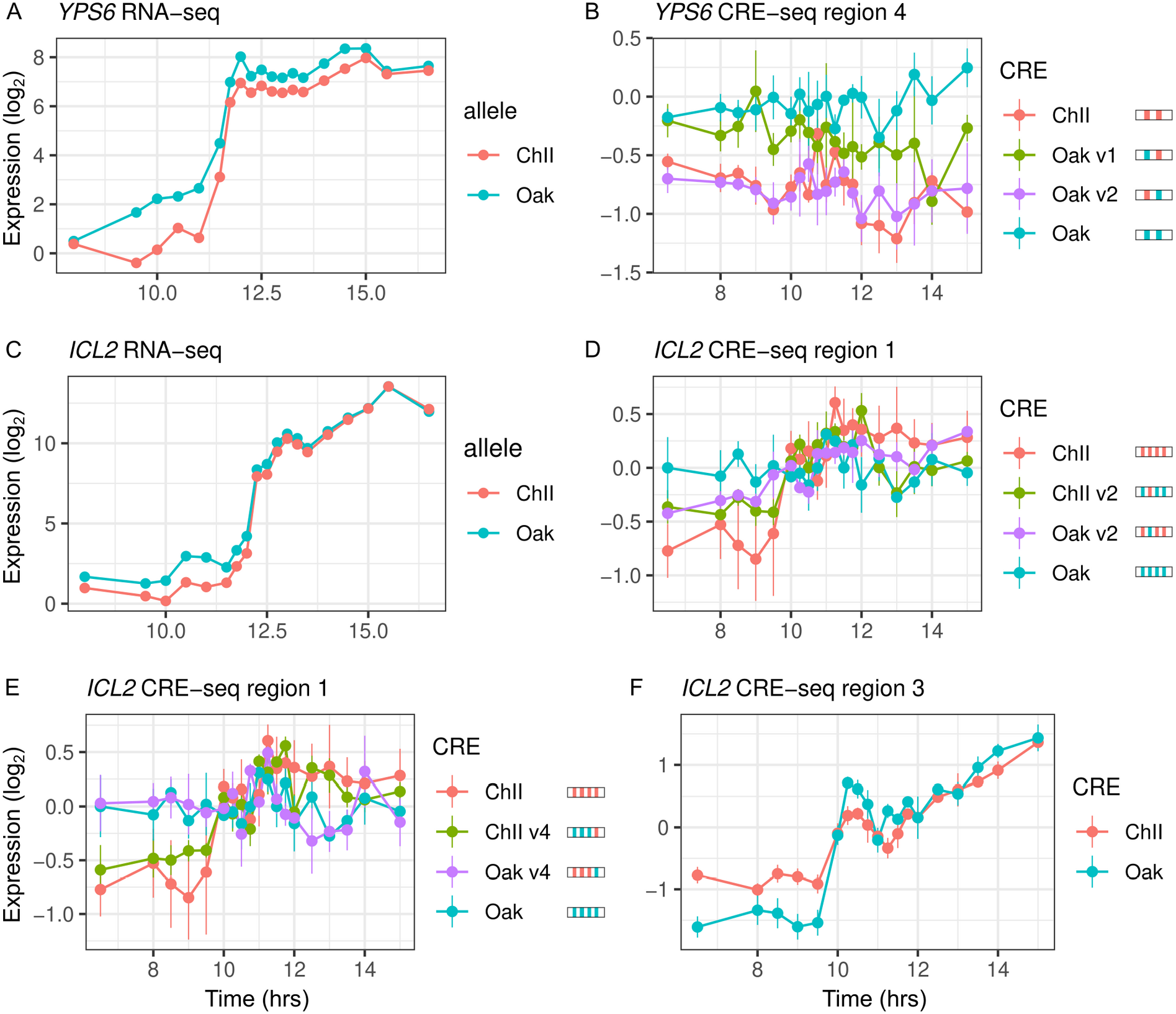
Intra-specific CREs show differences in expression levels and dynamics. (A) *YPS6* shows allele differences in endogenous expression levels and dynamics. (B) CRE region 4 of the *YPS6* promoter shows allele differences in expression levels, but is not correlated with endogenous expression patterns. Substituting the Oak InDei into the ChII allele (Oak v1) increases expression levels but substituting the Oak SNP into the ChII allele (Oak v2) has no effect. (C) *ICL2* shows allele differences in endogenous expression dynamics. Of the four SNPs and one InDei that differentiate the region 1 CRE alleles, two SNPs (v2 and v4) alter expression dynamics in both the Oak and ChII background (D and E). (F) CRE region 3 of *ICL2* has a single InDei between the Oak and ChII alleles and also shows allele differences in expression dynamics. For panels B, D and E, CRE alleles are shown by rectangles with colored ticks to indicate the Oak and ChII variants. Bars indicate standard errors.

*Cis*-regulatory variants that affect expression levels have been associated with conserved promoter regions and disruption of transcription factor binding sites (Renganaath et al., 2020). We found no difference in PhastCons conservation scores or change in binding site scores between the 35 variants associated with expression dynamics and those that were not associated (Figure S7). Because the number of variants is small we also examined whether PhastCons scores or binding site scores improved the genome-wide logistic regression. Neither PhastCons or binding site scores improved the association between upstream SNPs and InDels and ASE dynamics. However, PhastCons scores did improve the significance of the association between upstream SNPs and ASE levels (Table S6).

### Inter-specific *cis*-regulatory variation

We also designed a CRE-seq library to test promoter divergence of 98 genes that exhibited ASE levels and/or dynamics in the *S. cerevisiae* × *S. uvarum* hybrid. We used the same design of five overlapping 130 bp CRE sequences covering 250 bp upstream of the transcription start site. There was an average of 32 substitutions and 4.0 gaps per region. Because there were too many differences to test individually, we generated chimeric CRE sequences containing either the first or second half of the sequence from the *S. cerevisiae* allele and the remaining half from the *S. uvarum* allele (Figure S5). The total library contained 1,808 CREs with four barcode replicates per CRE.

We again tested whether the synthetic promoter regions could recapitulate expression of the endogenous genes and whether there were differences in expression levels or dynamics between the *S. cerevisiae* and *S. uvarum* alleles. We measure CRE expression during the diauxic shift in the same hybrid background used to measure RNA-seq. Out of 452 regions, 220 were correlated with RNA-seq expression (FDR < 0.05) and 28 genes had no region correlated. Similar to intra-specific comparisons, more regions showed differences in expression dynamics (113) than expression levels (2) between the two species’ alleles (Table 2).

Under an additive model, expression driven by chimeric sequences should lie within the range of the two parental species and can be used to map parental differences to the proximal or distal portion of the *cis*-regulatory region (Figure S8). However, chimera expression may also lie outside of the parental range as a consequence of binding site turnover or other interactions between divergence in the proximal and distal promoter regions. Such *cis*-regulatory interactions are thought to be common (Zheng et al., 2011), and do not require expression divergence between the parental species.

To map expression divergence and identify chimeras outside of the parental range, we tested each of the two chimeras for differences with each parent. Out of the 113 regions with parental species differences in expression dynamics, 57 were consistent with the proximal and 14 were consistent with the distal region explaining the difference. For example, the proximal region of *SDH4* can explain the entire difference between the parental species’ alleles (Figure 5A and 5D). Examining all the regions (n=348), 88 chimeras showed expression dynamics that differed from both parents (FDR < 0.05). The majority of these chimeras are not intermediate between the two parents; in 63 cases the expression distance between the two parents was less than the average distance of the chimera to either parent, and in 56 cases there was no difference between the two parents. As examples, *MDM36* shows chimera expression between the two parents (Figure 5B and 5E), and *IDP2* shows chimera expression outside the parental range (Figure 5C and 5F). For the 63 chimeras with high expression distance from both parents, the expression of the chimera was outside the two parental values for most (21.6/27) of the time-points.

**Figure 5.**
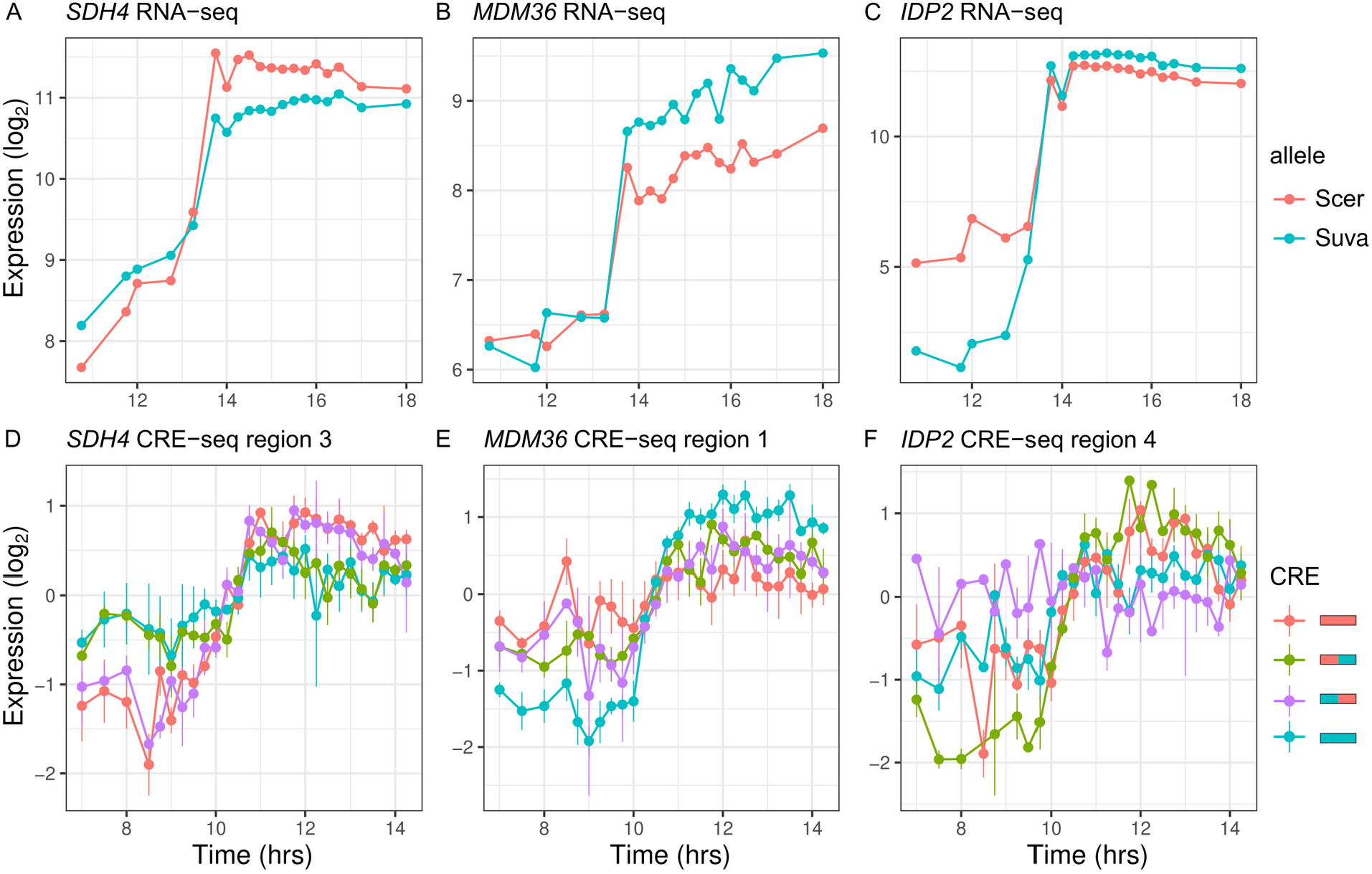
Inter-specific CREs show differences in expression dynamics. (A-C) Endogenous expression of *S. cerevisiae* (Scer) and *S. uvarum* (Suva) alleles of *SDH4*, *MDM36* and *IDP2*. (D-F) CRE-seq expression of region 3 (*SDH4*), region 1 (*MDM36*) and region 4 (*IDP2*) for parental S. *cerevisiae* (red) and S. *uvarum* (blue) CRE alleles and both chimeric CREs. *SDH4* shows expression divergence maps to the proximal promoter region, *MDM36* and *IDP2* show chimera expression that differ from both parents, with the *MDM36* chimeras being between the two parents and *IDP2* chimeras being outside the two parents. Bars indicate standard errors.

## Discussion

*Cis*-regulatory sequence control the activation and repression of genes in response to cellular and environmental signals, and the consequences of variation within these sequences are relevant to our understanding of evolution and human disease. Much progress has been made in identifying *cis*-regulatory variants responsible for changes in gene expression levels. In this study, we identified and characterized *cis*-acting variation in gene expression dynamics. We find that while gene expression dynamics are interrelated to expression levels, they differ in the types and identities of variants perturbing them. Below we discuss the relationship between gene expression dynamics and other aspects of gene regulation and how this fits into our understanding of regulatory evolution.

### Gene expression dynamics are an integrated component of *cis*-regulatory variation

Despite the potential importance of changes in gene expression dynamics, they are not easily disentangled from other aspects of gene expression such as levels and noise. In our data roughly one third of genes that exhibit ASE dynamics also exihibit ASE levels (Table 1). This overlap is not unexpected as the tests for ASE dynamics and levels are not mutually exclusive. For example, genes that exhibit differences in expression levels after but not before the diauxic shift by definition also exhibit differences in expression dynamics during the diauxic shift. While less common, we do find genes where ASE differences are greatest during the diauxic shift, consistent with a change in the rate or timing of gene activation/repression (Figure 1). How can the rate or timing of expression be altered without affecting levels? Prior work has shown that chromatin mutants can slow gene activation without impacting levels (Barbaric et al., 2001; Floer et al., 2010). While this does not exclude the possibility of changes in binding sites, it points to interaction between transcription factor binding and chromatin in mediating expression dynamics.

Gene expression dynamics may also be interrelated to stochastic noise in expression. During the diauxic shift there is cell to cell heterogeneity in gene expression and this leaky regulation can give the appearance of a slower rate of activation at the population level (New et al., 2014; Venturelli et al., 2015). Put differently, an increase is noise is expected to blur a sharp transition between activated and repressed states. However, noise, levels and dynamics are also not entirely dependent on one another. For example, gene expression often occurs in bursts with expression levels more dependent on burst size, noise more dependent on burst frequency (Cai et al., 2006; Carey et al., 2013), and timing dependent on both changes in burst size and frequency. Thus, the covariation between expression levels, dynamics and noise likely shapes how gene expression evolves within and between species.

### *Cis*-regulatory variants underyling expression dynamics

Previous studies have characterized *cis*-regulatory variants underlying gene expression levels (Doniger et al., 2008; Patwardhan et al., 2009; Gymrek et al., 2016; Tewhey et al., 2016; van Arensbergen et al., 2019; Hill et al., 2020; Renganaath et al., 2020). We find that *cis*-regulatory variants underlying expression dynamics are similar but not identical to those that affect expression levels.

Based on the genome-wide logistic regression we found InDels have larger effects than SNPs and stronger associations with variation in promoter regions compared to coding or 3’ regions. The larger effect of InDels is consistent with an eQTL mapping study in *Drosophila* (Massouras et al., 2012), and the stronger association with promoter regions is consistent with an eQTL mapping study in yeast (Kita et al., 2017). However, a limitation of the genome-wide associations is that the number of SNPs and InDels is correlated among 5’, coding and 3’ regions. Thus, the strongest associations are most likely to be causal but weaker associations could result from correlations with other regions. For the association with 3’ variants it is worth noting that ASE may be driven by variants that affect mRNA decay rather than transcription (Cheng et al., 2017).

We also found 5’ InDels have stronger associations with ASE dynamics compare to ASE levels. However, we did not find an over-representation of InDels among causal variants identified in the CRE-seq assay. This difference may be a consequence of a small sample size (n=35). But it could also reflect our selection of genes with the largest differences in ASE dynamics to test and promoters with multiple SNPs being more likely to cause large ASE differences than those with multiple InDels, which are more rare.

A previous study of *cis*-regulatory variants that affect expression levels in yeast found multiple *cis*-regulatory variants per gene, and that *cis*-regulatory variants are more likely to disrupt conserved sequences and alter trancription factor binding sites (Renganaath et al., 2020). We found that nearly half of the genes (8/20) had more than one variant (2-5) associated with ASE dynamics, but *cis*-regulatory variants associated with ASE dynamics were not associated with conserved sequences or changes in predicted transcription factor binding sites. Beyond technical differences, these differences could be related to differences in *cis*-regulatory variants underlying ASE levels versus dynamics or to the strains used in each study. Strain differences may be relevant since we used variants between two wild strains Oak and ChII, whereas Renganaath *et al*. (2020) used a wine and laboratory strain, the later of which has evolved under relaxed selection and has more deleterious variants (Gu et al., 2005; Doniger et al., 2008). Consistent with the possibility of differences between *cis*-regulatory variants that affect ASE levels versus dynamics, we found that SNP conservation scores improved the significance of genome-wide logistic regression for ASE levels but not dynamics (Table S6).

Although powerfull in throughput, the CRE-seq reporter assay has a number of limitations relevant to our results. First, 130 bp CRE sequences do not capture the entire promoter and different regions of a promoter often generate different patterns of expression. For example, a variant that modulates expression may have little or no effect unless upstream activation sequences are also included in the CRE. However, it is also possible that a variant affects expression regardless of the presence or absence of other elements. The extent to which the effects of individual binding sites are dependent on other sites forms the basis for the difference between the enhanceosome and billboard models of *cis*-regulatory sequences (Arnosti and Kulkarni, 2005). A second limitation of our study is that high-throughput reporter assays perform better with high levels of replication. Prior high-throughput reporter assays have used tens or hundreds of barcode replicates per allele (Tewhey et al., 2016; Renganaath et al., 2020). Our use of only four barcode replicates per allele likely limited our ability to detect variants that affect ASE levels. This limitation applies less to ASE dynamics which are unaffected by the mean expression of any single barcoded CRE. Given these limitations, not all *cis*-regulatory variants assayed by CRE-seq may have been detected.

### Evolution of *cis*-regulatory sequences

Chimeric *cis*-regulatory sequence from different species often show loss of function and support the binding site turnover model, whereby the chance gain of a redundant binding site enables loss of another site without adverse effects on expression (Ludwig et al., 2000, 2005; Arnold et al., 2014). We find chimeric promoter regions from *S. cerevisiae* and *S. uvarum* often generate expression dynamics outside of the parental species’ range. The chimeras thus provide evidence for constraints on gene expression dynamics and indicate that there are frequently non-additive (epistatic) interactions between substitutions that occur in different lineages and promoter regions. However, we also find that *cis*-regulatory variants within *S. cerevisiae* are not greatly enriched at conserved sites. These two observations are consistent with a neutral model of expression divergence (Fay and Wittkopp, 2008), whereby small changes in dynamics within species are neutral, but when neutral changes in different lineages are brought together they yield expression patterns that lie outside of the parent range and are unlikely to be tolerated within a species. It is also possible that constraints on expression dynamics depend on expression levels or noise. Indeed, the fitness effects of noise depend on expression levels (Duveau et al., 2018). This emphasizes the importance of characterizing how fitness altering *cis*-regulatory variants affect all aspects of gene expression to understand the evolution of *cis*-regulatory sequences.

## Methods and Materials

### Strains

Three strains of *S. cerevisiae*, one of *S. paradoxus* and one of *S. uvarum* were crossed to a common reference YJF153, a derivative of an *S. cerevisiae* oak isolate from North America (Table S1). The three intra-specific hybrids were generated through crosses to a wine strain from North America (UCD2120 × YJF153 = YJF1460) and two wild strains from China (HN6 × YJF153 = YJF1454 and SX6 × YJF153 = YJF1455). Strains were chosen to reflect a range of divergence. The two strains from China are from two of the most divergent *S. cerevisiae* lineages: China I (HN6) and China II (SX6)(Wang et al., 2012). The inter-specific hybrids were generated using *S. paradoxus* (N17 × YJF153 = YJF1453) and *S. uvarum* (CBS7001 × YJF153 = YJF1484). Diploid hybrids were generated by mixing strains of opposite mating type and selecting for dominant drug resistance markers present at the *HO* locus.

### RNA-sequencing time-course

Each hybrid strain was cultured in 125 mL YPD at 250 rpm at 30°C with an initial density of ~3×10^6^ cells/mL. Approximately ~3×10^8^ cells were taken at each time-point, centrifuged, supernatant removed and flash frozen in liquid nitrogen. A total of 19 samples were collected during the switch from fermentation to respiration with the most intense sampling occurring every 15 minutes after glucose depletion (Figure S1). Glucose depletion was measured using a Glucose (GO) Assay Kit (Sigma-Aldrich). RNA was extracted with phenolchloroform and mRNA purified by oligo-dT (Dynabeads mRNA Direct kit, Invitrogen). cDNA libraries were made by reverse transcription, fragmentation and adaptor ligation by Washington University’s Genome Technology Access Center (GTAC). Adaptors contained 7 bp indexes for multiplexing samples. The pooled equimolar libraries of 95 samples (19 sampling time × 5 strains) were paired-end sequenced (2 × 40 bp) using three runs of an Illumina NextSeq. A total of 1,255.9 million paired-end reads were generated.

### Reference genomes and variants

The genomes of the three *S. cerevisiae* strains were sequenced to generate reference genomes for mapping RNA-sequencing reads and to identify variants associated with allele-specific expression (ASE). DNA was extracted (YeaStar DNA kit, Zymo Research), libraries were generated by GTAC and paired end (2 × 101 bp) reads were generated using an Illumina HiSeq 2500 resulting in 5.2 to 5.9 million paired reads per strain. Reads were mapped to the S288c reference genome (R64-1-1) using BWA v0.7.5, (Li, 2013) and duplicates were marked using PicardTools v1.114 (http://broadinstitute.github.io/picard/). Single nucleotide polymorphisms (SNPs) and insertions/deletions (InDels) were called using GATK’s HaplotypeCaller v3.3-0 (Van der Auwera et al., 2013) following InDel realignment, base recalibration and variant recalibration using known SNPs and InDels. Known SNPs (21,327) and InDels (4,748) were identified using GATK after BWA mapping of 27 assembled genomes (Table S7) to the S288c reference genome. Variants were filtered to remove variable sites with calls in fewer than 20 strains and minor allele frequencies of less than 15%. We considered these “known” variants since they were identified from an independent set of strains assembled with both Sanger and 454 sequencing reads, and were found present in multiple strains. Using a tranche filter of 99.9 derived from the known variant call set we identified 222,589 SNPs and 20,485 InDels within the three *S. cerevisiae* strains and S288c. The Oak strain is most closely related to the Wine strain (76,659 variants), followed by China II (111,591 variants), China I (128,346 variants).

Variants were used to generate reference genomes for mapping RNA sequencing reads. Reference genomes were generated using the S288c genome as a template and GATK’s FastaAlternateReferenceMaker command to incorporate variants present in each strain. Genome annotations were generated using liftOver (Hinrichs et al., 2006) to transfer S288c annotations to each of the three other genomes. For mapping inter-specific hybrid reads we used reference genomes and annotations (Scannell et al. 2011) for *S. cerevisiae* (S288c), *S. paradoxus* (N17) and *S. uvarum* (CBS7001).

### Gene expression measurements

RNA sequencing reads were mapped to combined reference genomes containing the genomes of both parental strains used to generate the hybrid in order to avoid mapping bias. Reads were demultiplexed using the Fastx-toolkit (http://hannonlab.cshl.edu/fastx_toolkit/) and then mapped to combined reference genomes using Bowtie2 v2.1.0 (Langmead and Salzberg, 2012) using the local alignment setting (--local) and a maximum of one mismatch in the seed alignment (-N 1). Duplicate reads were marked using PicardTools. Htseq-count (Anders et al., 2015) was used to quantify allele-specific expression (ASE) by counting reads that mapped to each allele in the combined reference genomes. Reads mapping to overlapping features (-mode union) and reads with a mapping quality less than 10 (default) were not counted. Thus, reads that mapped equivalently to both alleles of a gene and assigned mapping quality of 0 or 1 by Bowtie2 were removed. For interspecific hybrids we used previous definitions of orthologous genes (Scannell et al., 2011). The fraction of mappable reads used for ASE was 81.58%, 82.62%, 83.11%, for the intra-specific hybrids YJF1454, YJF1455 and YJF1460 and 86.16%, 86.57% for the inter-specific hybrids YJF1450 and YJF1484, respectively. None of the intra-specific hybrids showed any mapping bias, except in YJF1454 genes on chromosome XIII R showed uniformly higher expression of the Oak compared to the ChI allele consistent with aneuploidy (Figure S2). The fraction of reads mapping to the common reference YJF153 was 52.58% (50.59%, after removing chrXIII R), 50.11%, 49.78%, 49.11%, 46.90% for YJF1454, YJF1455, YJF1460, YJF1453 and YJF1484. The median number of allele-specific reads per time-point was 2.9 million reads (range: 1.5-4.8) for intra-specific hybrids and 2.0 million reads (range: 1.4-3.3) for inter-specific hybrids.

### Statistical analysis of differentially expressed alleles

Allele-specific expression counts were normalized using DESeq’s blind method (Love et al., 2014). Differences in expression levels were tested using a weighted linear model: *f_i_*=0.5+*β*_0_*+e*, where *f_i_* is the normalized frequency of the YJF153 reference allele over time-point *i*, *β_0_* is the deviation from 0.5, and *e* is the error with weights based on the total number of reads at each time-point. Differences in expression dynamics were tested using a weighted Durbin-Watson test for an autocorrelation across timepoints of the allele differences. The weighted Durbin-Watson test is based on the *lmtest* package of R where the residuals (*e*) from the same linear model used above are used to calculate the test statistic:

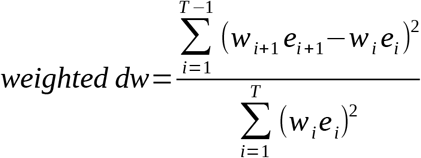

where, *w_i_* and *e_i_* are the weights (read counts) and residuals for sample *i* out of *T* time-points. For both tests we used false discovery rate (FDR) cutoffs of 0.01.

Genes with low read counts were removed from the analysis: either those with less than an average of 20 reads per time-point or more than seven time-points with no reads. This filter eliminated an average of 692 genes per hybrid, and left a total of 4,703 genes with data in all five hybrids.

### Gene expression clusters

Kmeans clustering was used to group genes by their combined allele expression profile and by their allele imbalance profile. For the combined gene expression profiles, 19,633 genes from all five hybrids with a significant (FDR < 0.01) autocorrelation in the combined (both alleles) profile over time were normalized (centered and scaled) and clustered into 12 groups. For the allele imbalance profiles, allele frequencies were centered but not scaled for genes with significant ASE dynamics (n=6,135) and clustered into 12 groups. For both, 336 genes without any reads at one or more time-points were removed since Kmeans clustering does not handle missing data.

### Association with variants

The numbers of SNPs and InDels for intra-specific hybrids were obtained from variant calling, and the corresponding counts for inter-specific hybrids were obtained from multiple sequence alignments without correction for multiple hits (Scannell et al., 2011). Upstream intergenic regions were defined by sequences between adjacent coding regions, except for cases of short (< 5 bp) or overlapping coding sequences where we extended the region to the next upstream gene. Downstream intergenic regions were defined as 80 bp downstream of the stop codon. Variant counts were obtained for all of the 4,703 genes with expression data except for 11 genes which only had counts for the intra-specific hybrids.

For logistic regression we predicted genes with differences in ASE levels and dynamics using: *logit*(*p*)=*b*_0_+*b*_1_*x*+*e*, where *p* indicates whether a gene shows significant ASE or not, *b_1_* is the regression coefficient, *x* is the predictive variable and *e* is the error. We used a Bonferroni cutoff for significance to correct for the 60 different regressions based on: 6 predictive variables: number of SNPs and InDels within upstream, downstream and coding regions, and 10 response variables: ASE dynamics and ASE levels in each of the 5 hybrids.

### CRE-seq

A high-throughput reporter assay (CRE-seq) was used to measure the activity of *cis*-regulatory elements by sequencing (Mogno et al., 2013). In this assay, a pool of synthetic *cis-*regulatory sequences are cloned en masse, YFP is inserted between the *cis*-regulatory element and the barcode, and the reporter library is then integrated into the genome. The activity of each CRE is measured by the ratio of barcode sequencing reads from an RNA relative to a DNA library of the pooled transformants. As described below, we used this reporter assay to measure expression of 7,268 and 7,232 synthetic CRE sequences, representing intra-specific and interspecific allele differences upstream of 69 and 98 genes, respectively.

### Synthesis

CREs were synthesized as part of a library of 200 bp oligos (Agilent). The synthetic oligos include a forward primer, RS1, CRE, RS2, RS3, RS4, BC, RS5, reverse primer, where CRE is the *cis*-regulatory element, RS1-5 are restriction sites and BC is a barcode (Figure S5). The barcodes were random 10bp sequences with a minimum of 2 differences and no restriction sites. The selected barcodes were checked to ensure no base composition bias. Target CRE sequences were defined by the 250 bp region upstream of the transcription start site (TSS) (Venters et al., 2011). For each target CRE, five 130 bp sliding windows of the sequence were generated using a 30 bp step size. Any 130 bp regions containing coding sequence of the next upstream gene were removed. Four replicates with different barcodes were generated for each CRE. CREs were generated separately for each strain allele unless the CRE window was identical between the strains. Transcription start sites (Xu et al., 2009) were found by liftOver from the reference (S288c) genome to the coordinates in the strain of interest. In cases where the CREs contained InDels, extra sequence was added to the gapped allele between RS2 and RS4 to keep the oligo length constant. CREs were also synthesized for intra-specific variants between the Oak (YJF153) and China II strain (SX6). These CREs were designed by replacing a single Oak variant by the China II variant and vice versa (Figure S5). Inter-specific chimeric CREs were generated by recombining the *S. cerevisiae* allele with *S. uvarum* allele at the center of each CRE (Figure S5). The intra-specific and inter-specific libraries included 7,268 and 7,232 synthetic CRE sequences representing 69 and 98 genes, respectively. Genes were chosen based on: the absence of restriction sites within the target CRE, an annotated TSS, significant ASE levels and/or ASE dynamics and inspection of the allele-specific expression differences. For the 69 genes used for the intra-specific libraries, 19 showed ASE levels, 10 showed ASE dynamics and 40 showed both. For the 98 genes used in the inter-specific library, 23 showed ASE level, 10 showed ASE dynamics and 65 showed both.

### Cloning

CRE-seq libraries were amplified (8 separate reactions of 10-cycle amplification), digested, gel purified and ligated into pIM202 (Mogno et al., 2013). This differs from the original protocol that used higher resolution acrylamide gel extraction since sequencing the amplified library showed only a small portion (<1%) of sequences were outside of the 200+/-5 bp resolution of the acrylamide gel. Ligated products were transformed into bacteria using electroporation and 80-100k colonies were pooled and the plasmid library was extracted. YFP along with 69 bp of the *TSA1* core promoter (Mogno et al., 2013) was inserted between the CRE and the barcode and then transformed back into bacteria by electroporation. At each step, PCR was used to ensure a low frequency of empty vector.

### Transformation into yeast

The reporter gene (YFP) with synthetic CREs and corresponding barcodes were integrated at the *URA3* locus (Figure S5 D). BstBI digested plasmids were transformed into the Oak strain (YJF186) using the LiAc method (Gietz et al., 1995). We collected ~100k colonies on complete minimal plates without uracil and integration was confirmed by PCR for 20/20 randomly selected colonies. The yeast strains transformed with intra-specific library were co-cultured with SX6 (YJF1375) in YPD at 30C overnight to allow mating and HygB and dsdA was used to select diploid cells. The pool of yeast strains transformed with inter-specific library were co-cultured with the *S. uvarum* strain (YJF1450) in YPD overnight at 30C to allow mating. Double selection with nourseothricin and dsdA was used to select diploid cells.

### Full length promoters

Three genes, *ALD5*, *GND2* and *PHO3*, were chosen to compare short and full length promoter constructs. Full length promoters from the Oak (YJF186) and ChII (SX6) strains were amplified from the TSS to the next upstream coding sequence. Four barcodes per construct and restriction sites were incorporated into the primers used for PCR. Each full length promoter with barcodes was cloned into the pIM202 plasmid and YFP was inserted between the full length promoter and the barcode. BstBI digested plasmids were transformed into the Oak strain at the *URA3* locus. Each of the transformed strains were mated with SX6 to form diploids.

### CRE-seq expression measurements

Pooled libraries were shaken at 250 rpm in 125 mL YPD at 30C with an initial density of 3×10^6^ cells/mL. Samples of 3×10^8^ cells were taken for DNA and RNA extraction between 6 hrs and 15 hrs, which spans the switch from fermentation to respiration. A total of 19 samples were taken for RNA measurements for the intra-specific library and a total of 27 samples were taken for the inter-specific library. For both libraries, sampling corresponded to the RNA-sequencing time-points but with more dense sampling for the interspecific library. For each library, samples were taken for DNA measurements at the first and last time-points. Cells were centrifuged for 30 seconds at 1000g and the pellets were immediately frozen with liquid nitrogen.

The abundance of each CRE in the library was measured by sequencing the barcodes from DNA and the expression of each CRE was measured by sequencing the barcodes from RNA extracts. DNA was extracted from the first and the last time-points using YeaStar Genomic DNA Kit (Zymo Research). RNA was extracted by YeaStar RNA Kit (Zymo Research) and residual DNA digested with DNase (Promega). mRNA was purified by oligo-dT (Invitrogen) and cDNA was made using SuperScript II Reverse Transcriptase (Invitrogen). mRNA was removed after first strand DNA synthesized with RNase H (New England BioLabs). Combinations of indexed Ion Torrent primers were used to amplify barcodes in the library for each time-point (Figure S5 D) using Phusion High-Fidelity PCR (New England BioLabs). To avoid sampling bias during PCR amplification, each sample was amplified with 4 PCR reactions and 20 cycles and then pooled. PCR pools were gel-extracted and cleaned individually, then pooled together at equivalent concentrations.

After demultiplexing the samples using the indexed Ion Torrent primer pairs, perfect matches to the CRE barcodes were found for 401 and 97 million reads from the intra- and inter-specific libraries. For one sample, three technical replicate libraries was generated (PCR and sequencing). The technical replicates showed an average correlation of 0.96 for barcode counts and were subsequently combined. We removed CREs that had either zero reads in more than a third of the RNA time-point samples or those with less than 100 reads on average in either the RNA or DNA samples. This filter left 6,820 intra-specific and 5,013 intra-specific CREs with an average of 16.4 and 3.2 million reads per time-point and a median of 1,278 and 357 reads per barcode across all time-points for the intra- and inter-specific libraries, respectively. The read count distribution of intra- and inter-specific libraries were normalized using DESeq2’s blind method (Love et al., 2014). Expression of each CRE allele was measured by the ratio of RNA to DNA counts. Barcodes with zero counts were treated as missing data in statistical models.

### Identification of significant expression differences

CRE-seq regions were tested for correlations (Pearson) with RNA-seq using the average expression of all barcodes for a given CRE region. For inter-specific CRE-seq, correlations were measured between the 19 timepoints closest to the RNA-seq time-points. CREs with significant differences in expression levels and dynamics were tested using the weighted linear model and weighted Durbin-Watson test, respectively. When testing for differences in expression levels using the linear model we used the average expression across all time-points for each barcode rather than treating each timepoint as an independent measure of expression levels. We chose this more conservative test to avoid cases where one barcoded CRE had substantially higher or lower expression across all time-points compared to the other replicate barcodes. Consequently, the power to detect differences in average CRE expression levels based on four barcode replicates was reduced. The test for expression dynamics was not affected by this problem since expression dynamics was measured by changes in the ratio of expression levels over time.

### Identification of variant and chimera differences

For the intra-specific library, we tested individual variants for those CREs with more than one difference between the Oak and ChII alleles. We used the same weighted tests for differences in expression levels and dynamics, but tested each variant genotype separately (Figure S8). For the inter-specific library we used the chimeric CREs to map differences to the proximal or distal part of each region showing significant differences between the parental *S. cerevisiae* and *S. uvarum* alleles. The left (distal) and right (proximal) promoter regions were separately tested using the genotype of the left and right portions of the CRE, respectively (Figure S8). Parental differences were classified as mapping to the proximal region if the proximal but not the distal genotype was significant. Distal mapping was similarly classified. Chimeric CREs that showed significant differences from both parents were also identified, and subsequently classified as outside or inside the parental range if their average expression distance (Euclidean) to each parent was greater or less than the distance between the two parents, respectively. A flow diagram of the number of CREs tested along with genotypes tests for significant variants and chimeras is presented in Figure S8.

### Associations with variant annotations

Variants were annotated with PhastCons and transcription factor binding site scores. Phastcons scores (Siepel et al., 2005) were obtain for yeast from the USCS genome browser. Scores were extracted for each SNP and the average score was obtained for InDels based on the two sites flanking the InDel and any scores within the InDel. Transcription factor binding motifs were obtained for 196 factors (Spivak and Stormo, 2012). For each variant we extracted 30 bp of sequence from the Oak and ChII genome on either side of the variant. We used Patser (Hertz and Stormo, 1999) to scan each sequence with each motif model and the best hit was recorded. A background nulceotide frequency of 36% GC was used and scores less than zero (equivalent likelihood between the motif and background model) were set to zero. The difference in score between the two alleles was calculated for each motif model and the maximum difference across all motif models was used as the binding site annotation score for each variant. PhastCons scores and binding site scores were each tested for association with CRE variants that were positive (n=35), negative (n=35), and all other intergenic variants (n=44,514) by ANOVA.

## Acknowledgements

We thank Feng-Yan Bai for sharing yeast strains and members of the Fay lab for their suggestions and comments.

## Data availability

Genome sequencing and assembly data were deposited into NCBI, see Table S1 and Table S7 for accessions. RNA sequencing data were deposited into NCBI’s GEO database under GSE165594. Analysis scripts, data and files underlying figures are available at https://doi.org/10.17605/OSF.IO/Y5748

## Supporting Information

Table S1. Strains used in this study.

Table S2. Kmeans clusters of gene expression dynamics.

Table S3. Kmeans clusters of allelic differences in expression.

Table S4. Logistic regression of ASE dynamics and levels.

Table S5. Average number of SNP and InDel differences within hybrids.

Table S6. Logistic regression with binding site and conservation scores.

Table S7. Genome assemblies used to identify known variants.

Figure S1. Sampling scheme for gene expression dynamics during the diauxic shift.

Figure S2. Chromosome 13R aneuploidy in YJM1454 (Oak × ChI).

Figure S3. Characteristics of differentially expression genes.

Figure S4. Gene expression dynamics.

Figure S5. Design of synthetic sequences for CRE-Seq.

Figure S6. Short CREs recapitulate longer CRE expression.

Figure S7. Binding site and conservation scores of variants.

Figure S8. Identification of significant differences for the intra-specific and inter-specific CRE-seq libraries.

