## Supplementary Figures for "*Cis*-regulatory variants affect gene expression dynamics in yeast"

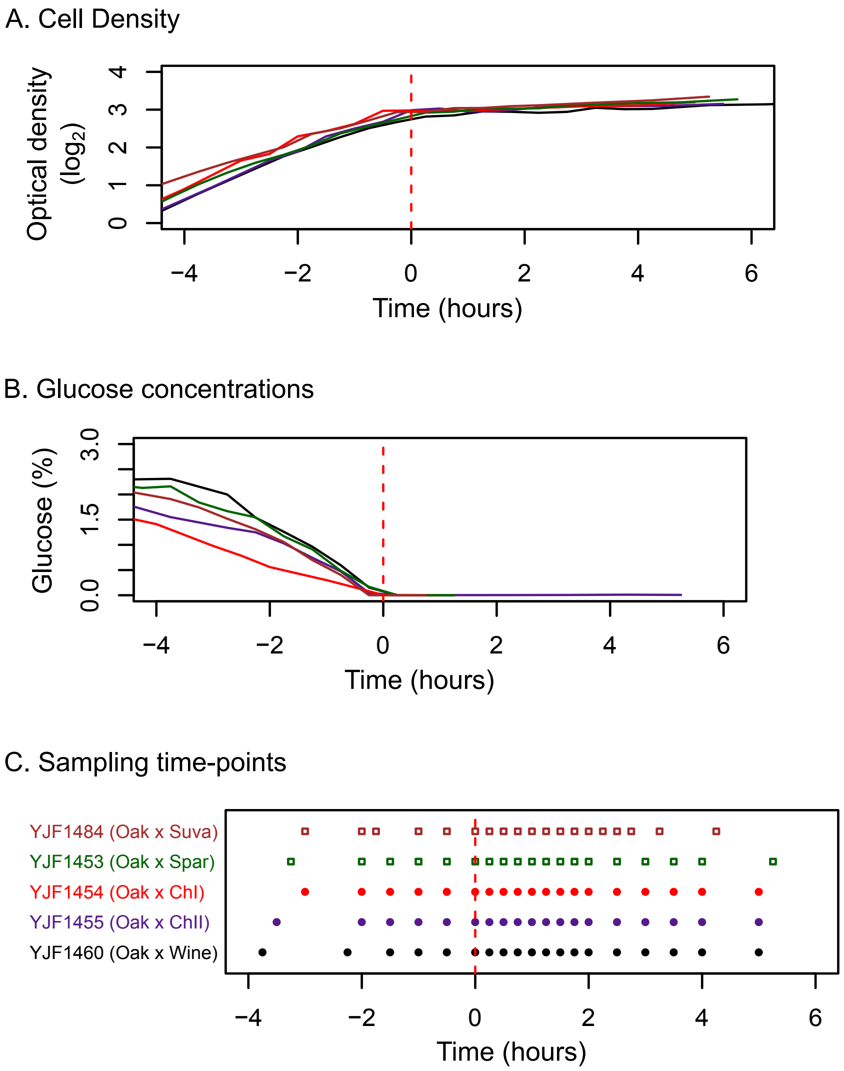


Figure S1. Sampling scheme for gene expression dynamics during the diauxic shift. Cell density (A) and glucose concentrations (B) were used to choose sampling time-points for gene expression measurements (C). Time of glucose depletion is marked as 0 hours by a red dashed line. Sample time-points were taken every 15 minutes for two hours after glucose depletion. Intra-specific hybrid samples are circles, inter-specific hybrid samples are squares.

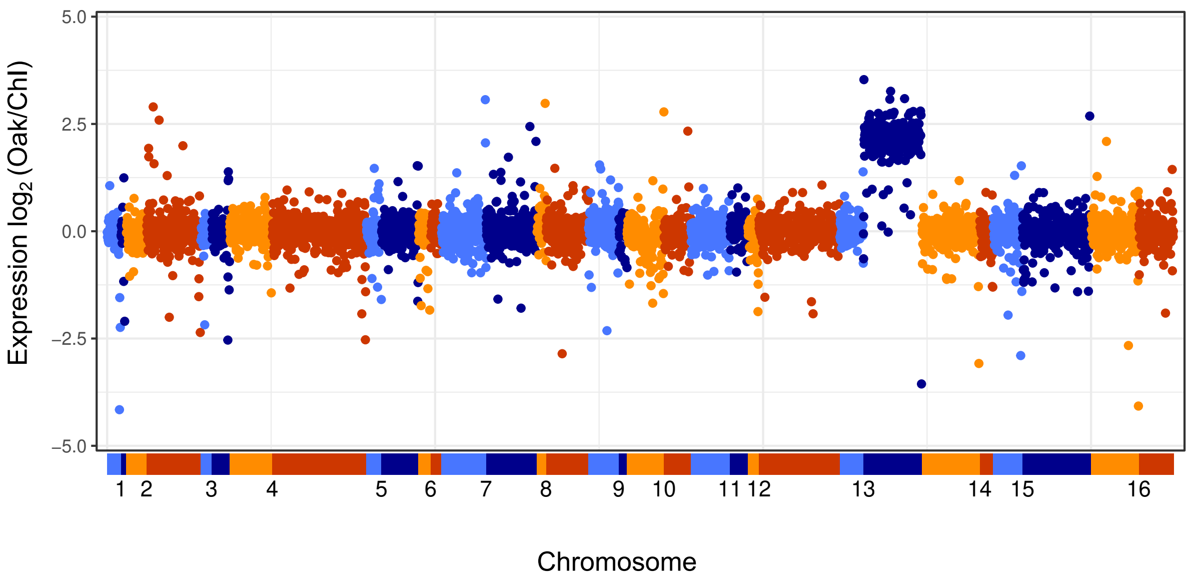


Figure S2. Chromosome 13R aneuploidy in YJM1454 (Oak x Chl). Expression of the Oak relative to the Chl allele for all genes ordered by position along each chromosome, where expression is the average across all time-points. Points are color coded by chromosome (alternating blue and orange, bottom), with light colors for the left arm and dark colors for the right arm.

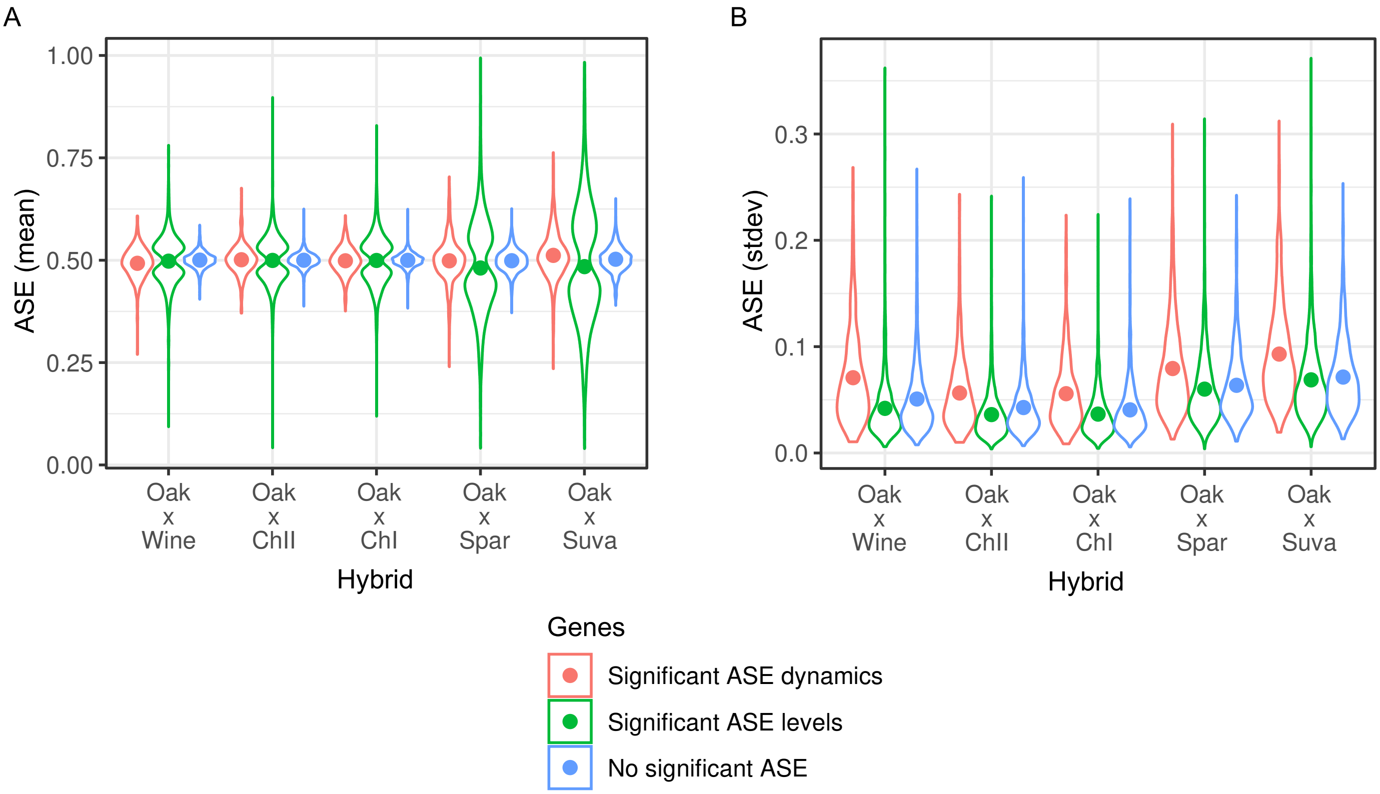


Figure S3. Characteristics of differentially expression genes. (A) Average ASE expression levels, and (B) standard deviation in ASE expression over time, for genes with significant ASE dynamics (red), ASE levels (green), and no ASE (blue) for each of the five hybrids. Points show the mean and density is indicated by the width of the color coded regions.

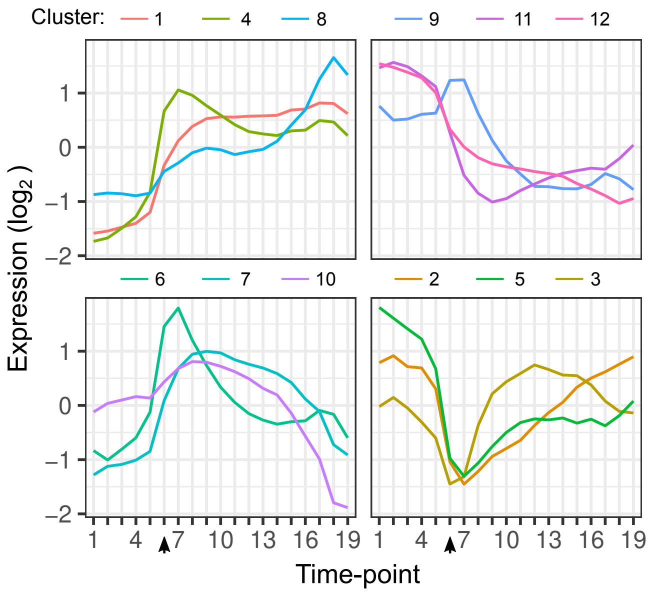


Figure S4. Gene expression dynamics. Each line shows the average expression of genes in each k-means cluster over time-points. Clustering is based on the expression of 4,703 genes from each hybrid. The arrow indicates the time-point when glucose was depleted.

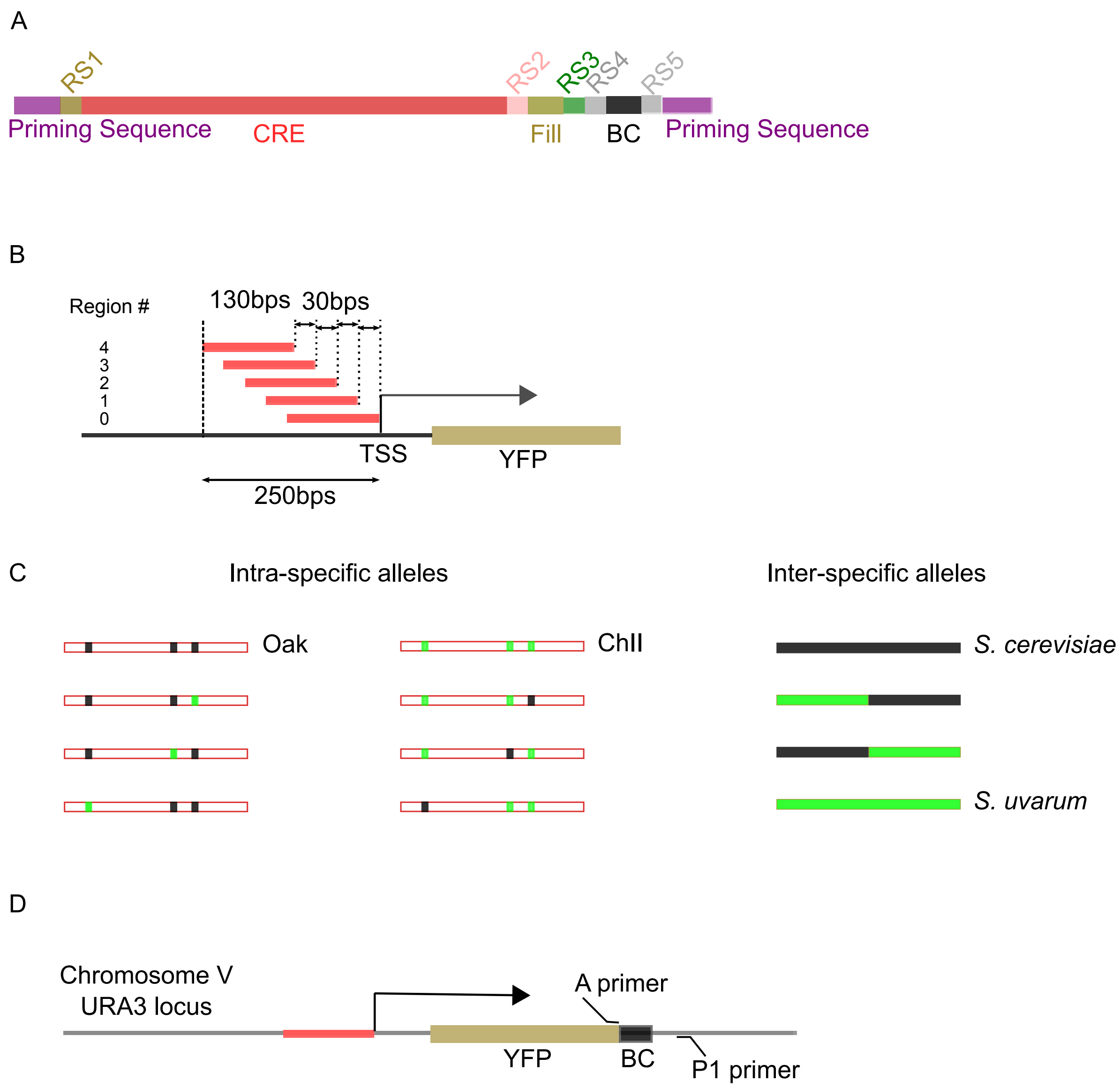


Figure S5. Design of synthetic sequences for CRE-Seq. (A) Synthetic CREs contain priming sequence to amplify, restriction sites (RS) for cloning, fill sequences to compensate different lengths due to any InDels, barcodes (BC), and CREs. (B) The designed 130 bp CREs with 30 bp steps covers 250 bp upstream of the TSS. (C) Intra-specific CREs include the Oak (YJF153, black) and ChII (SX6, green) alleles, as well as each variant substituted into the Oak and ChII background. Inter-specific CREs include the *S. cerevisiae* (black) and *S. uvarum* (green) alleles as well as two chimeric alleles designed with recombination in the center of CREs. (D) The reporter gene (YFP) with the barcoded CREs were integrated at the *URA3* locus. The A and P1 primers were used for barcode sequencing of extracted RNA and DNA.

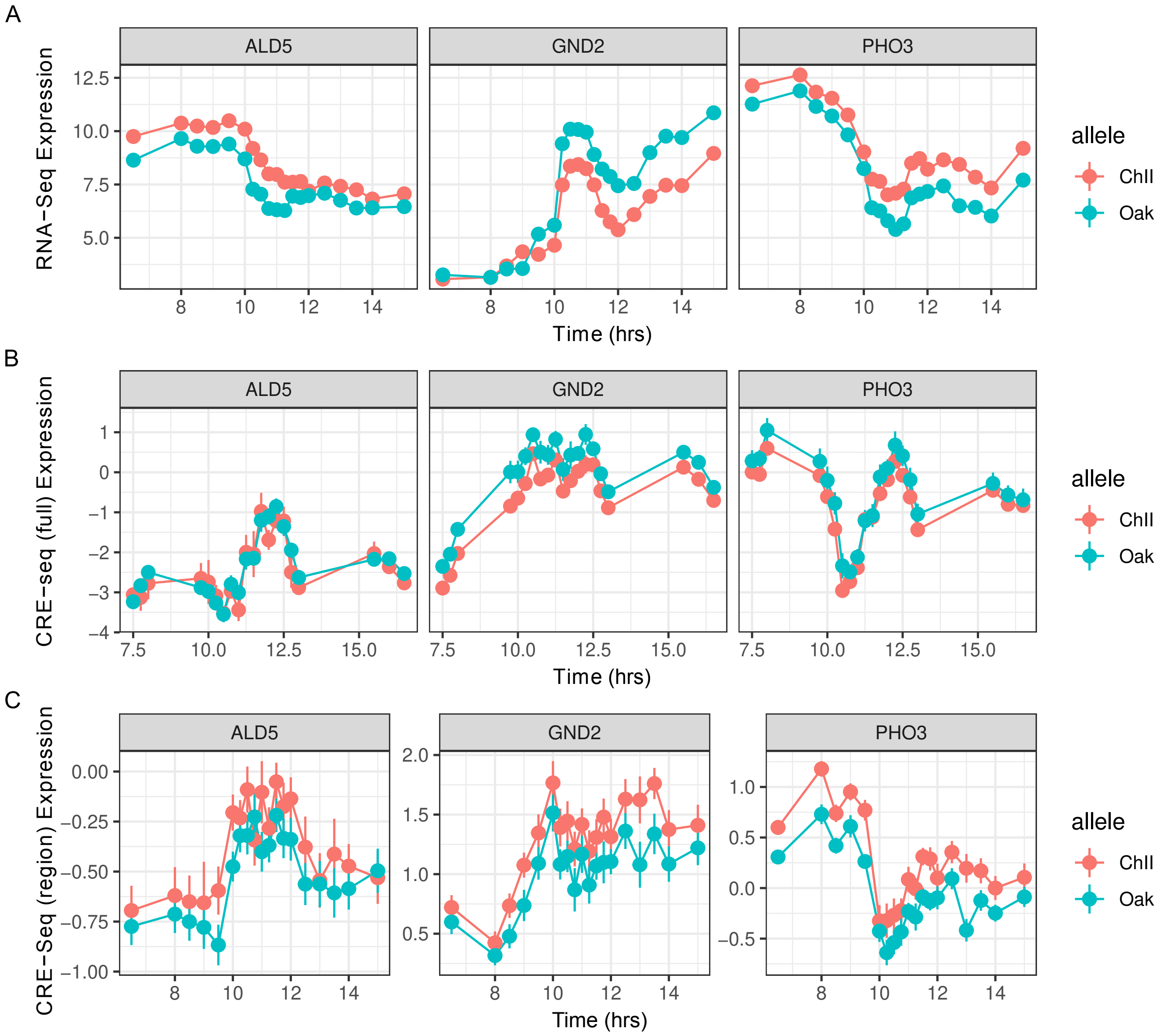


Figure S6. Short CREs recapitulate longer CRE expression. Gene expression dynamics of three genes (*ALD5*, *GND2*, *PHO3*) measured by RNA-seq (A) in comparison to CRE-seq expression from full length promoters (B) and CRE-seq expression from short 130 bp promoter regions (C). Points and lines show the mean and standard error from replicate barcodes of the Oak and ChII alleles. Expression is shown on a log2 scale.

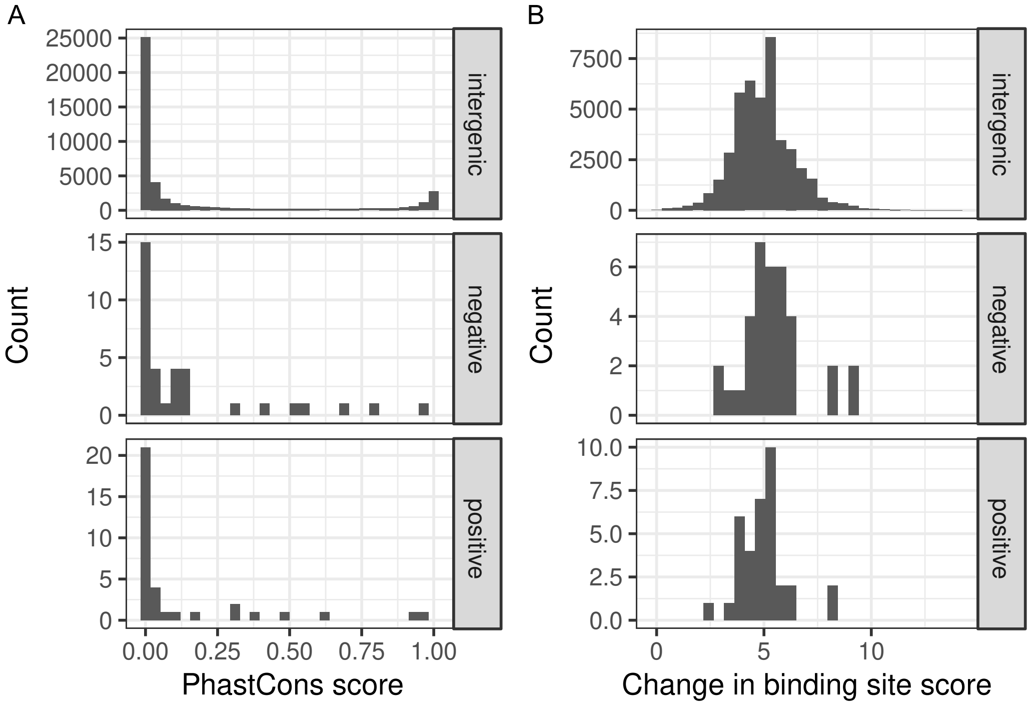


Figure S7. Binding site and conservation scores of variants. (A) Histograms of PhastCons conservation scores (0 least conserved, 1 most conserved). (B) Histograms of the maximum change in transcription factor binding scores cause by a variant across 196 binding site models. The distribution is shown for variants associated with CRE-expression dynamics (positive, n=35) or not (negative, n=35), and all other intergenic variants in the Oak x ChII hybrid (n=44,514).

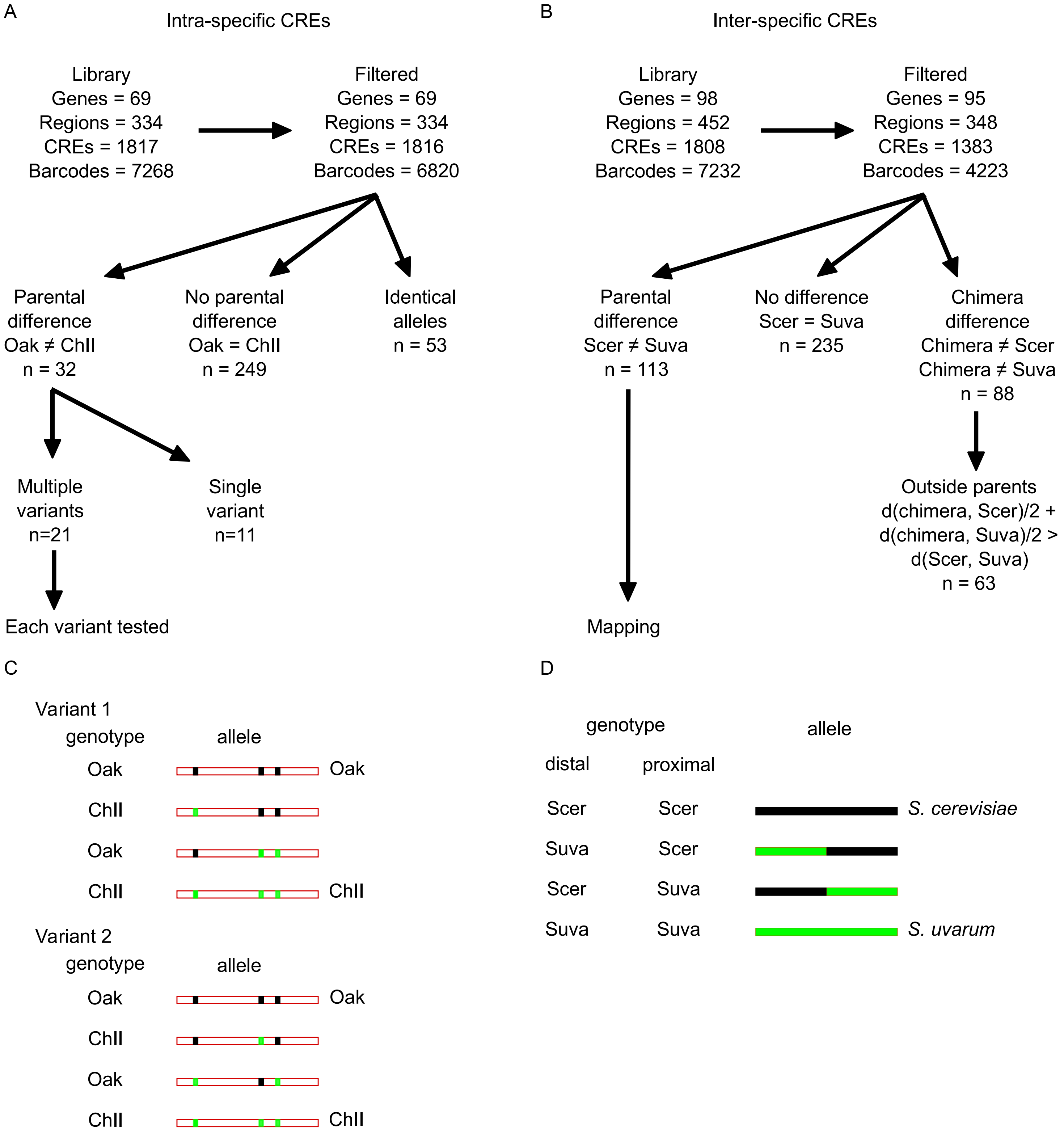


Figure S8. Identification of significant differences for the intra-specific (left) and inter-specific (right) CRE-seq libraries. The number of genes, promoter regions, CREs (including variants and chimeras) and barcodes is shown for the intra-specific (A) and inter-specific (B) library. The same numbers are shown after filtering to remove low abundance barcodes and CREs without replicates. For the intra-specific data, differences between the parental (Oak and ChII) alleles were identified for all CRE regions that were not identical between Oak and ChII. Variants within these CRE region were tested when there was more than one difference between the Oak and ChII alleles. Examples of how two variants were tested is shown by the variant genotype and the allele background in which it occurs (C). For the inter-specific library, differences between the parental *S. cerevisiae* (Scer) and *S. uvarum* (Suva) alleles were identified, and subsequently mapped using the genotype of the proximal or distal region of the parental and chimeric alleles (D). Chimeras with expression outside the parental range were identified by those where a chimera differed from both parental alleles and where the average distance of the chimera to each parent is greater than the distance between the two parents.
